# Cross-reactive coronavirus antibodies with diverse epitope specificities and extra-neutralization functions

**DOI:** 10.1101/2020.12.20.414748

**Authors:** Andrea R. Shiakolas, Kevin J. Kramer, Daniel Wrapp, Simone I. Richardson, Alexandra Schäfer, Steven Wall, Nianshuang Wang, Katarzyna Janowska, Kelsey A. Pilewski, Rohit Venkat, Rob Parks, Nelia P. Manamela, Nagarajan Raju, Emilee Friedman Fechter, Clinton M. Holt, Naveenchandra Suryadevara, Rita E. Chen, David R. Martinez, Rachel S. Nargi, Rachel E. Sutton, Julie E. Ledgerwood, Barney S. Graham, Michael S. Diamond, Barton F. Haynes, Priyamvada Acharya, Robert H. Carnahan, James E. Crowe, Ralph S. Baric, Lynn Morris, Jason S. McLellan, Ivelin S. Georgiev

## Abstract

The continual emergence of novel coronavirus (CoV) strains, like SARS-CoV-2, highlights the critical need for broadly reactive therapeutics and vaccines against this family of viruses. Coronavirus spike (S) proteins share common structural motifs that could be vulnerable to cross-reactive antibody responses. To study this phenomenon in human coronavirus infection, we applied a high-throughput sequencing method called LIBRA-seq (Linking B cell receptor to antigen specificity through sequencing) to a SARS-CoV-1 convalescent donor sample. We identified and characterized a panel of six monoclonal antibodies that cross-reacted with S proteins from the highly pathogenic SARS-CoV-1 and SARS-CoV-2 and demonstrated a spectrum of reactivity against other coronaviruses. Epitope mapping revealed that these antibodies recognized multiple epitopes on SARS-CoV-2 S, including the receptor binding domain (RBD), N-terminal domain (NTD), and S2 subunit. Functional characterization demonstrated that the antibodies mediated a variety of Fc effector functions *in vitro* and mitigated pathological burden *in vivo*. The identification of cross-reactive epitopes recognized by functional antibodies expands the repertoire of targets for pan-coronavirus vaccine design strategies that may be useful for preventing potential future coronavirus outbreaks.

## INTRODUCTION

The emergence of a novel coronavirus (CoV) SARS-CoV-2, the causative agent of COVID-19, has resulted in a worldwide pandemic, threatening the lives of billions and imposing an immense burden on healthcare systems and the global economy. SARS-CoV-2, the seventh coronavirus known to infect humans, is a member of the *Betacoronavirus* genus which includes the highly pathogenic SARS-CoV-1 and MERS-CoV, as well as endemic variants OC43-CoV and HKU1-CoV^1^. Recent coronavirus outbreaks and the threat of future emerging zoonotic strains highlight the need for broadly applicable coronavirus therapeutic interventions and vaccine design approaches^2^.

Coronaviruses utilize the homotrimeric Spike (S) protein to engage with cell-surface receptors and enter host cells. S consists of two functional subunits: S1 and S2. S1 facilitates attachment to target cells and is composed of the N-terminal domain (NTD) and the receptor-binding domain (RBD), whereas S2, which encodes the fusion peptide and heptad repeats, promotes viral fusion^3,4^. To facilitate cell entry, human coronaviruses employ different host factors; however, SARS-CoV-1 and SARS-CoV-2 both utilize the cell-surface receptor, angiotensin converting enzyme 2 (ACE2)^5^. Additionally, SARS-CoV-2 S shares 76% amino acid identity with SARS-CoV-1 S^1^. Furthermore, S serves as a dominant antibody target and is a focus of countermeasure development for the treatment and prevention of COVID-19 infection^6,7^. S proteins from the *Betacoronavirus* genus share multiple regions of structural homology and thus could serve as targets for a cross-reactive antibody response^8^. Identifying cross-reactive antibody epitopes can inform rational design strategies for vaccines and therapies that target multiple highly pathogenic coronaviruses, which will be of value both for the current and potential future outbreaks.

Numerous potent neutralizing antibodies against SARS-CoV-2 have been discovered, including multiple candidates currently in clinical trials for prophylactic and acute treatment of COVID-19^9–13^. With the goal of achieving cross-neutralization, investigation of SARS-CoV-2/SARS-CoV-1 cross-reactive antibodies has focused primarily on the RBD epitope. This has resulted in the identification of a number of SARS-CoV-2/SARS-CoV-1 cross-reactive antibody candidates^12,14,15^. However, the diversity of epitopes and functions beyond virus neutralization have not been extensively explored for cross-reactive antibodies^16–18^. Evidence of Fc effector function contributing to protection *in vivo* against SARS-CoV-1^19^ and SARS-CoV-2^20^ suggests that the role of antibodies beyond neutralization (“extra-neutralization” functions) may be a crucial component of protection and an important consideration in vaccine design strategies for coronaviruses^17,21–23^. Defining the genetic features, epitope targets, and Fc effector functions of cross-reactive antibodies can provide insights into current therapeutic strategies and may provide alternative approaches for the prevention and treatment of coronavirus infection.

In this study, we investigated antibody cross-reactivity across the *Betacoronavirus* genus at monoclonal resolution. To do this, we applied LIBRA-seq (Linking B Cell receptor to antigen specificity through sequencing), a recently developed high-throughput antibody screening technology that allows for determination of B cell receptor sequence and antigen reactivity simultaneously for many single B cells^24^. From a convalescent SARS-CoV-1 donor sample, we identified and characterized SARS-CoV-2/SARS-CoV-1 cross-reactive human antibodies that target multiple, distinct structural domains of S, mediate Fc effector functions, and mitigate pathological burden *in vivo*. A better understanding of the epitope specificities and functional characteristics of cross-reactive coronavirus antibodies may translate into strategies for current vaccine design efforts and additional measures to counteract potential future pandemic strains.

## RESULTS

### LIBRA-seq Applied to a SARS-CoV-1 Convalescent Donor

To identify cross-reactive antibodies to multiple coronavirus antigens, LIBRA-seq was applied to a PBMC sample from a donor previously infected with SARS-CoV-1 over ten years prior to sample collection. The antigen screening library consisted of eight oligo-tagged recombinant soluble antigens: six coronavirus trimer antigens (SARS-CoV-2 S, SARS-CoV-1 S, MERS-CoV S, MERS-CoV S1 (with foldon domain), OC43-CoV S, HKU1-CoV S) and two HIV trimer antigens from strains ZM197 and CZA97 as negative controls (**Figure 1A**). After the antigen screening library was mixed with donor PBMCs, antigen positive B cells were enriched by fluorescence activated cell sorting and processed for single-cell sequencing (**Supplemental Figure 1A**). After bioinformatic processing, we recovered 2625 cells with paired heavy/light chain sequences and antigen reactivity information (**Supplemental Figure 1B**). Overall, LIBRA-seq enabled rapid screening of PBMCs from a patient sample, with recovery of paired heavy/light chain sequences and antigen reactivity for thousands of single B cells.

**Figure 1.**
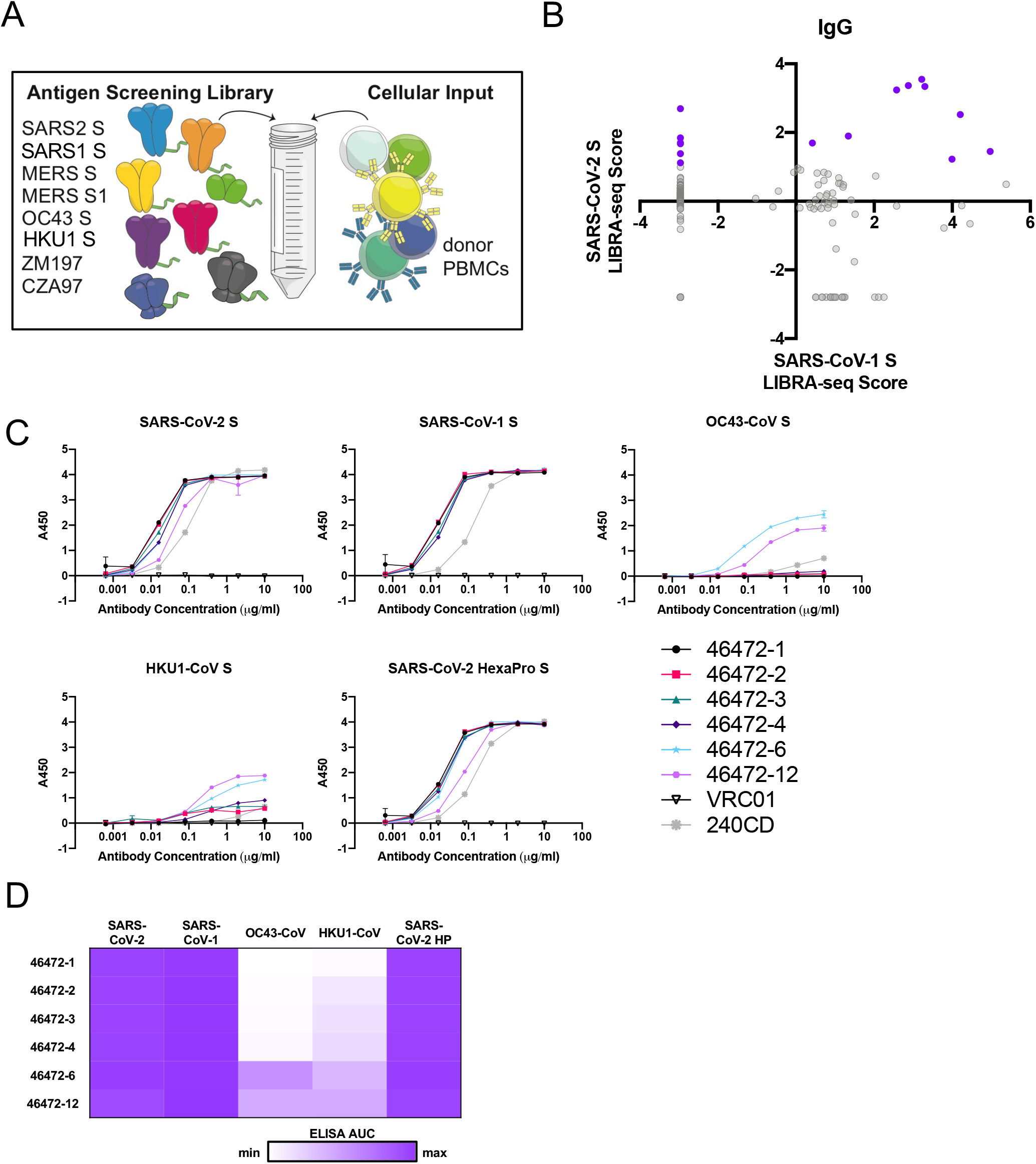
Identification of coronavirus cross-reactive antibodies from SARS-CoV-1 convalescent PBMC sample using LIBRA-seq. (**A**) Schematic of DNA-barcoded antigens used to probe a SARS-CoV-1 donor PBMC sample. The LIBRA-seq experiment setup consisted of eight oligo-labelled antigens in the screening library: SARS-CoV-2 S, SARS-CoV-1 S, MERS-CoV S, MERS-CoV S1, OC43-CoV S, HKU1-CoV S, and two HIV negative controls (ZM197, and CZA97). (**B**) LIBRA-seq scores for SARS-CoV-1 (x-axis) and SARS-CoV-2 (y-axis) for all IgG cells recovered from sequencing are shown as circles. The 15 lead antibody candidates are highlighted in purple. (**C**) Antibodies were tested for binding to SARS-CoV-2 S (S-2P), SARS-CoV-1 S (S-2P), OC43-CoV S (S-2P), HKU1-CoV S (S-2P), and SARS-CoV-2 S (HexaPro) by ELISA. HIV-specific antibody VRC01 is used as a negative control. Anti-SARS-CoV-1 mouse antibody 240CD was also used (BEI Resources). ELISAs were performed in technical duplicates with at least two biological duplicates. (**D**) ELISA binding data against the antigens are displayed as a heatmap of the AUC analysis calculated from the data in Figure 1C, with AUC of 0 displayed as white, and maximum AUC as purple. ELISAs were performed in technical duplicates with at least two biological duplicates.

### Identification of SARS-CoV-2 and SARS-CoV-1 Cross-reactive Antibodies

With a goal of identifying antibodies that were cross-reactive to multiple coronavirus S proteins, we prioritized lead candidates based on their sequence features and LIBRA-seq scores. We selected 15 antibody candidates that exhibited diverse sequence features and utilized a number of different variable genes for expression and characterization (**Figure 1B, Supplemental Figure 1C**). These antibodies displayed a broad range of somatic hypermutation levels (83-98%) and a variety of CDRH3 and CDRL3 lengths (6-24 and 5-12 amino acids, respectively) (**Supplemental Figure 1C**). Antibodies 46472-1, 46472-2, 46472-3, 46472-4, 46472-6, and 46472-12 showed binding to SARS-CoV-1 S and SARS-CoV-2 S by ELISA (**Figure 1C-D, Supplemental Figure 1D**). Further, antibodies 46472-6 and 46472-12 bound to S proteins from endemic OC43-CoV and HKU1-CoV, albeit generally at lower levels (**Figure 1C-D, Supplemental Figure 1D**). Although the six monoclonal antibodies showed reactivity by ELISA to the MERS antigen probe used in the LIBRA-seq screening library, antibody binding to other independent preparations of this protein was inconsistent, so we could not definitively confirm MERS S reactivity. Overall, the application of the LIBRA-seq technology enabled the identification of a panel of cross-reactive antibodies that recognize the S antigen from multiple coronaviruses.

### Cross-reactive Coronavirus Antibodies Target Diverse Epitopes on S

To elucidate the epitopes targeted by the cross-reactive antibodies, we performed binding assays to various structural domains of S as well as binding-competition experiments. First, we assessed antibody binding to the S1 and S2 subdomains of SARS-CoV-2. Antibodies 46472-1, 46472-2, 46472-3, and 46472-4 bound to the S2 domain, whereas 46472-6 and 46472-12 recognized the S1 domain but targeted different epitopes, the NTD and RBD, respectively (**Figure 2A-C, Supplemental Figure 2A-B**). Although 46472-12 bound to the RBD, it did not compete with ACE2 for binding to SARS-CoV-2 S (**Supplemental Figure 2C**). To determine whether the antibodies targeted overlapping or distinct epitopes, we performed competition ELISA experiments and found that the S2-directed antibodies 46472-1, 46472-2, and 46472-4 competed for binding to S (**Figure 2D**). This pattern was observed for both SARS-CoV-2 and SARS-CoV-1 S. Of note, this competition group did not include S2-directed antibody 46472-3, revealing the identification of multiple cross-reactive epitope targets on S2 (**Figure 2D**). Further, binding assays with glycan knockout mutants and mannose competition experiments revealed no notable glycan dependence for antibody reactivity to S (**Supplemental Figure 2D-E**). Lastly, we measured antibody autoreactivity, and found that with the exception of 46472-6 binding to Jo-1, none of the antibodies showed autoreactivity against the tested antigens (**Figure 2E**). Together, these data suggest that these cross-reactive antibodies are coronavirus-specific and target multiple, diverse epitopes on the S protein (**Figure 2F**).

**Figure 2.**
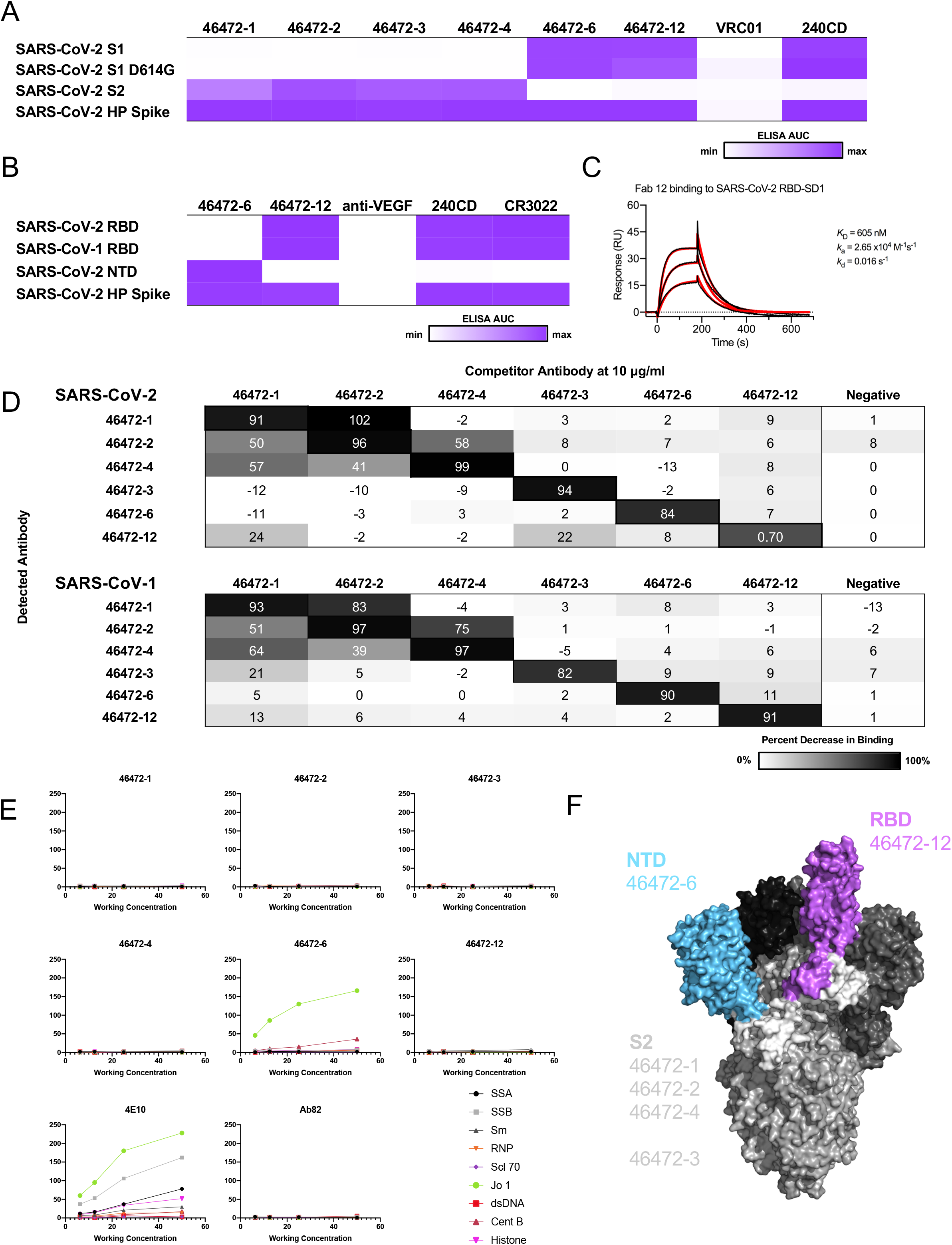
Epitope mapping of cross-reactive antibodies. (**A**) For cross-reactive coronavirus antibodies, ELISA binding data against the antigens are displayed as a heatmap of the AUC analysis calculated from the data in **Figure S2A** and (**B**) for SARS-CoV-2 S1 reactive antibodies, ELISA binding data against the RBD and NTD are displayed as a heatmap of the AUC analysis calculated from the data in **Figure S2B**. ELISA AUC is displayed as a heat map. AUC of 0 is displayed as white and maximum AUC as purple. ELISA data are representative of at least two independent experiments. Anti-HIV antibody VRC01 and anti-VEGF antibody are shown as a negative control and anti-SARS-CoV-1 antibody 240CD is shown as positive control. (**C**) Surface plasmon resonance binding of 46472-12 Fab to SARS-CoV-2 RBD. Affinity measurements are shown to the right of the graph. (**D**) Cross-reactive antibodies were used in a competition ELISA to determine if binding of one antibody affected binding of another. Competitor antibodies were added at 10 μg/ml, and then detected antibodies were added at 0.1 μg/ml. The percent reduction in binding compared to binding without a competitor is shown. An anti-HIV antibody was also used as a negative control. ELISAs were performed in technical duplicates with at least two biological duplicates. (**E**) Antibodies were tested for autoreactivity against a variety of antigens in the Luminex AtheNA assay. Anti-HIV antibody 4E10 was used as a positive control and Ab82 was used as a negative control. (**F**) Cross-reactive coronavirus antibodies target a variety of epitopes on the SARS-CoV-2 S protein, including the RBD, NTD, and S2 domains, highlighted on the structure (PDB: 6VSB). Antibodies targeting each epitope are listed and color coded for each domain.

### Functional Characterization of Cross-reactive Coronavirus Antibodies

Next, we characterized our cross-reactive antibody panel for functional activity. Although none of the antibodies neutralized SARS-CoV-1 or SARS-CoV-2 (**Supplemental Figure 3A-B**), all six antibodies demonstrated a range of Fc effector functions. Notably, all antibodies showed antibody-dependent cellular phagocytosis (ADCP) *in vitro* for SARS-CoV-2 S (**Figure 3A**). In particular, antibody 46472-12 which targets RBD showed greater ADCP activity compared to the other cross-reactive antibodies and the SARS-CoV-1/SARS-CoV-2 cross reactive RBD antibody control, CR3022^25^ (**Figure 3A, Supplemental Figure 3C**). Further, we tested and confirmed ADCP activity against SARS-CoV-1 for two antibodies, 46472-4 and 46472-12, illustrating that these antibodies can mediate antiviral function against multiple coronaviruses (**Figure 3B, Supplemental Figure 3D**). In a coated SARS-CoV-2 S assay (see Methods), the cross-reactive antibodies also mediated trogocytosis, an Fc-mediated immune function defined by the removal of cell membrane proteins from S-coated and opsonized cells to effector cells, which results in rapid cell death and antigen transfer^26^ (**Figure 3C, Supplemental Figure 3E**). Interestingly, only the S2-targeting antibodies in our panel (46472-1, 46472-2, 46472-3, and 46472-4) mediated trogocytosis for cell-surface expressed SARS-CoV-2 S (**Figure 3D, Supplemental Figure 3F**). Lastly, none of the antibodies promoted complement deposition (ADCD) (**Figure 3E, Supplemental Figure 3G**). Together, these results revealed that although this cross-reactive antibody panel is non-neutralizing, the six antibodies are capable of mediating a spectrum of Fc effector functions.

**Figure 3.**
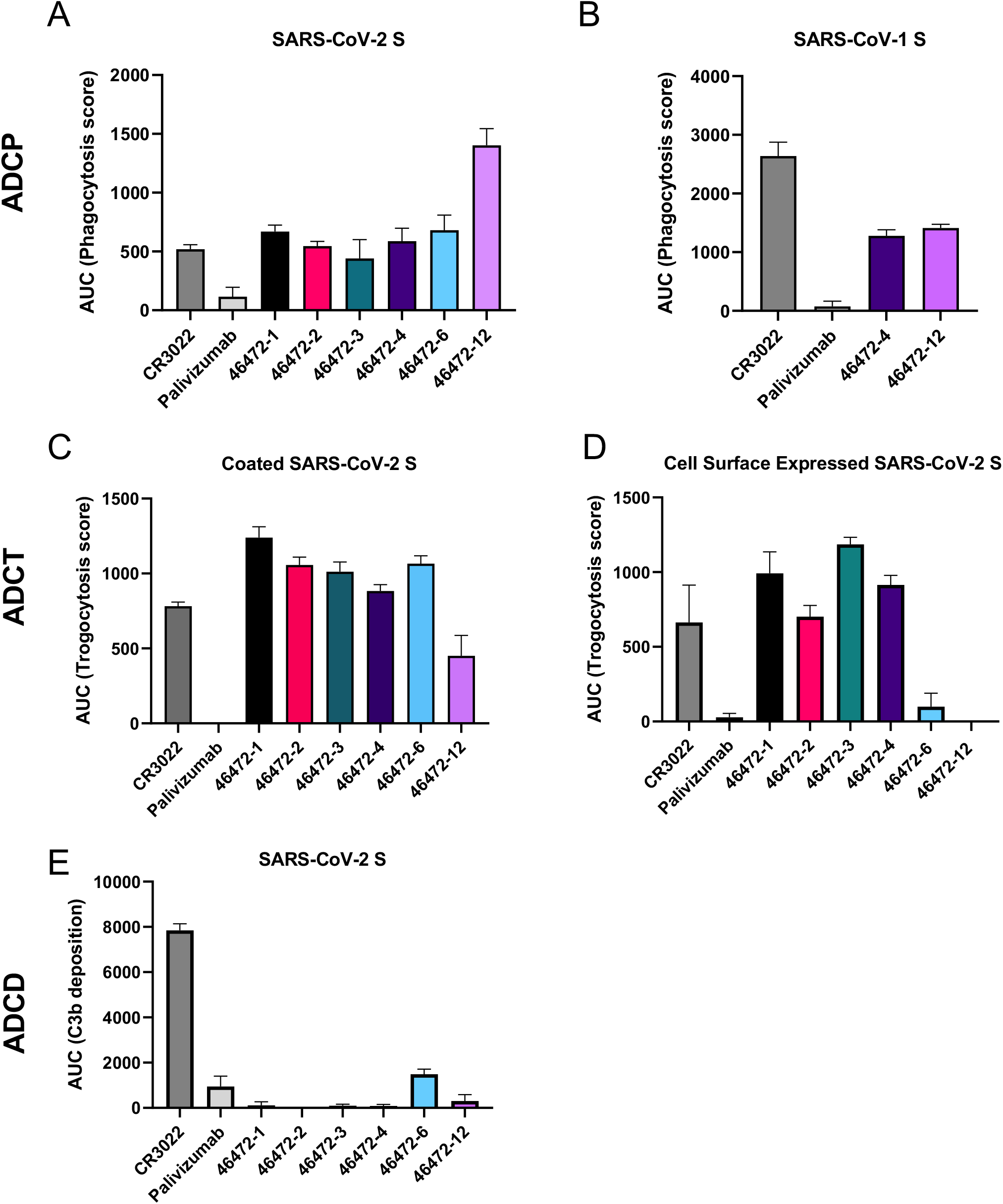
Functional activity of cross-reactive coronavirus antibodies. (**A**) Cross-reactive coronavirus antibodies were tested for antibody-dependent cellular phagocytosis activity (ADCP) against SARS-CoV-2 S, compared to positive control antibody CR3022 and negative control Palivizumab, an anti-RSV antibody. Area under the curve of the phagocytosis score is shown, calculated from data in **Figure S3C**. (**B**) 46472-4 and 46472-12 were tested for antibody-dependent cellular phagocytosis activity against SARS-CoV-1 S, compared to CR3022 antibody and anti-RSV antibody Palivizumab. Area under the curve of the phagocytosis score is shown, calculated from data in **Figure S3D**. (**C**) Cross-reactive coronavirus antibodies were tested for antibody-dependent cellular trogocytosis (ADCT) activity against SARS-CoV-2 S coated on cells, compared to positive control CR3022 and anti-RSV antibody Palivizumab. Area under the curve of the trogocytosis score is shown, calculated from data in **Figure S3E**. (**D**) Cross-reactive coronavirus antibodies were tested for antibody-dependent cellular trogocytosis activity against SARS-CoV-2 S displayed on transfected cells, compared to positive control CR3022 and anti-RSV antibody Palivizumab. Area under the curve of the trogocytosis score is shown, calculated from data in **Figure S3F**. (**E**) Cross-reactive coronavirus antibodies were tested for antibody-dependent complement deposition (ADCD) activity against SARS-CoV-2 S, compared to positive control CR3022 and anti-RSV antibody Palivizumab. Area under the curve of the C3b deposition score is shown, calculated from data in **Figure S3G**.

Since non-neutralizing SARS-CoV-2 antibodies with Fc effector function activity have not been extensively characterized *in vivo*, these results prompted us to test antibodies 46472-4 and 46472-12 for prophylaxis in a murine infection model using a mouse-adapted virus strain (SARS-CoV-2 MA)^27,28^(**Figure 4A**). Although there were no differences in survival and viral load between experimental and control groups, the hemorrhage scores (see Methods) for 46472-4 and 46472-12 were similar to positive control CR3022, and all three groups were lower than the scores for isotype control 2D22 (p<0.01, ordinary one-way ANOVA with multiple comparisons), suggesting a reduction in pathological burden (**Figure 4B, Supplemental Figure 4A-B**). To evaluate the *in vivo* effect of these antibodies in a more stringent challenge model in 12-month old BALB/c mice, we increased the viral dose from 1×10^3^ to 1×10^4^ PFU. In this experiment, mice that received antibody 46472-12 exhibited the best survival rate (4/5 at day 4), compared to all other treatment groups (including CR3022 as a positive control and DENV-2D22 as a negative control), although statistical significance was not achieved (**Figure 4C, Supplemental Figure 4C**). There were no significant differences in viral load between groups; however, the surviving animals from the 46472-4 and 46472-12 groups showed lower hemorrhagic pathology scores in harvested mouse lungs compared to the negative control treatment group (p<0.001, ordinary one-way ANOVA with multiple comparisons) (**Figure 4D, Supplemental Figure 4D**). Notably, animals treated with the positive control, CR3022, had higher hemorrhage scores than animals treated with 46472-4 and 46472-12 (p<0.001, ordinary one-way ANOVA with multiple comparisons), although the statistical analysis may be limited by the small numbers of surviving animals for some of the groups (**Figure 4D**). Together, the *in vivo* experiments suggest these cross-reactive antibodies could contribute to counteracting coronavirus infection in prophylaxis.

**Figure 4.**
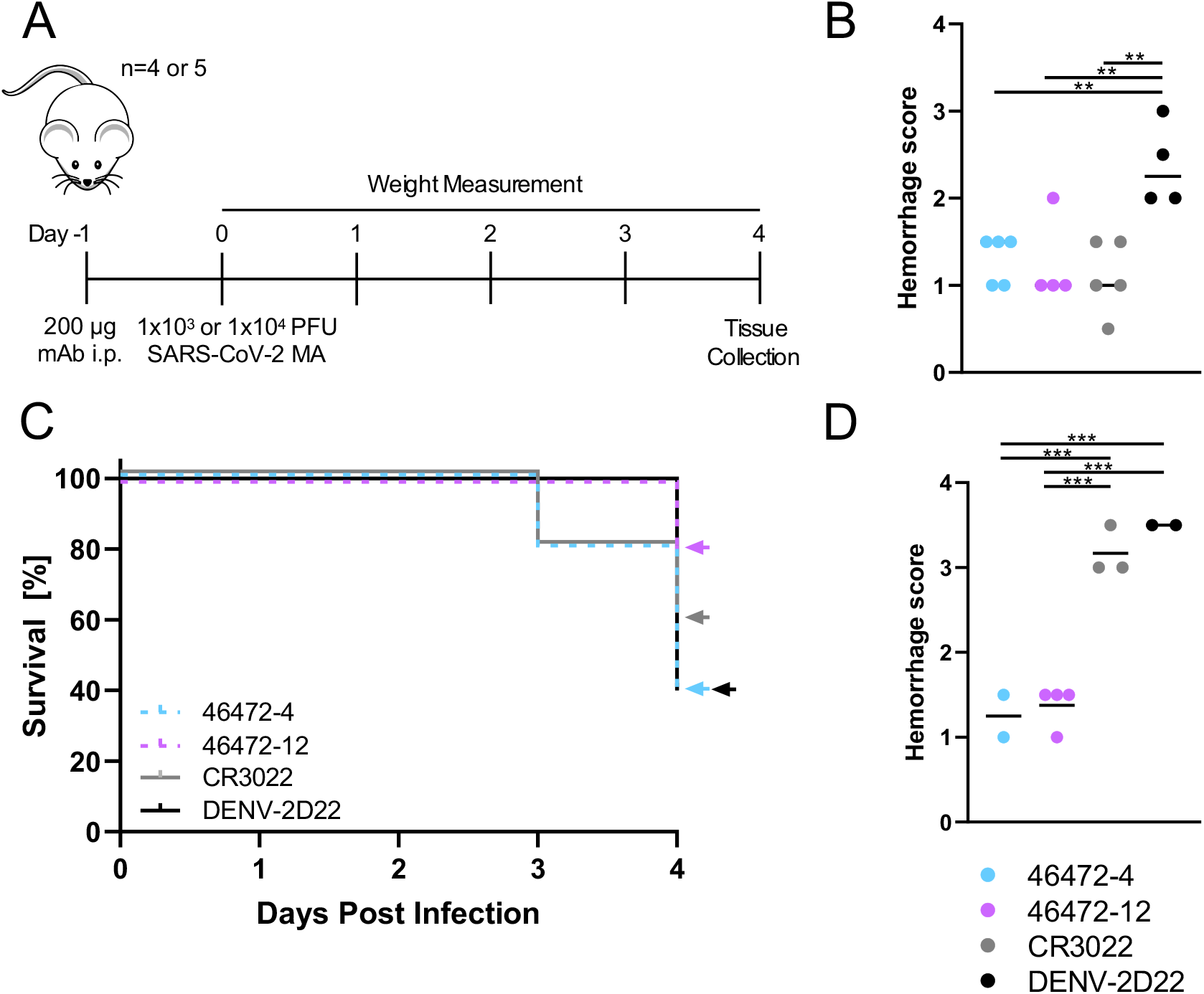
*In vivo* effects of cross-reactive antibodies. (**A**) Timeline of the prophylactic antibody experiment in SARS-CoV-2 mouse adapted (MA) in vivo infection model. 200 μg antibody was given via intraperitoneal route to 12-month old female BALB/c mice 12 hours prior to virus inoculation (n= 4 or 5 per group). 1×10^3^ or 1×10^4^ PFU infectious dose of SARS-CoV-2 MA was administered intranasally for the low dose and high dose experiments, respectively. Weights were measured daily, and on day 4 tissue was collected for histopathology and viral load quantification. (**B**) Lung hemorrhage scores of gross pathology are shown for each low dose (1×10^3^ PFU of SARS-CoV-2 MA) treatment group. An ordinary one-way ANOVA test with multiple comparisons was performed. (**C**) For the experiment treating with 1×10^4^ PFU of SARS-CoV-2 MA, percent survival for each antibody group is shown. 2/5, 4/5, 3/5, and 2/5 mice survived to day 4 for antibodies 46472-4, 46472-12, CR3022 and isotype control DENV-2D22 respectively. (D) Lung hemorrhage scores of gross pathology are shown for each high dose (1×10^4^ PFU of SARS-CoV-2 MA) treatment group. An ordinary one-way ANOVA test with multiple comparisons was performed.

## DISCUSSION

Here, we described a set of cross-reactive *Betacoronavirus* antibodies isolated from a convalescent SARS-CoV-1 donor. The antibodies targeted diverse epitopes on S, including the S2 subdomain as well as the RBD and NTD on S1, and were shown to be functional *in vitro*. Additionally, two of these antibodies were tested and demonstrated activity *in vivo*, displaying a reduction in lung hemorrhage score, while effects on viral load were not definitive. Although the precise *in vivo* effects of these antibodies have not been elucidated, the Fc effector function profiles in the absence of neutralization points to a potential role for extra-neutralization activity in the reduction of pathological burden. Evidence of protection associated with Fc effector function in SARS-CoV-1^19^, SARS-CoV-2^20,21^ (also described in Atyeo et al. 2020, available at SSRN: https://ssrn.com/abstract=3612156), and other infectious diseases including influenza, Ebola, and HIV, motivates further investigation into its contribution for the treatment of COVID-19^29-32^.

Given the ongoing SARS-CoV-2 pandemic and the potential for future zoonotic coronavirus pathogens to emerge, coronavirus vaccine and therapeutic development is of paramount importance^33–36^. Antibodies that can cross-react with multiple coronavirus variants are primary targets as potential broadly reactive therapies. Such antibodies can further reveal cross-reactive epitopes that will serve as templates for the development of broadly protective vaccines. Understanding the spectrum of cross-reactive epitopes targeted by human antibodies, as well as the functional role that such antibodies have in preventing and treating coronavirus infection, are therefore critical for medical countermeasure development. In particular, the identification of functional cross-reactive antibodies that target diverse epitopes on S will present a viable avenue for pan-coronavirus vaccine design strategies.

## Acknowledgements

We thank Angela Jones, Latha Raju, and Jamie Roberson of Vanderbilt Technologies for Advanced Genomics for their expertise regarding NGS and library preparation; David Flaherty and Brittany Matlock of the Vanderbilt Flow Cytometry Shared Resource for help with flow panel optimization; and members of the Georgiev laboratory for comments on the manuscript. The Vanderbilt VANTAGE Core provided technical assistance for this work. VANTAGE is supported in part by CTSA grant 5UL1 RR024975-03, the Vanderbilt Ingram Cancer Center (P30 CA68485), the Vanderbilt Vision Center (P30 EY08126), and NIH/NCRR (G20 RR030956). This work was conducted in part using the resources of the Advanced Computing Center for Research and Education at Vanderbilt University (Nashville, TN). Flow cytometry experiments were performed in the VUMC Flow Cytometry Shared Resource. The VUMC Flow Cytometry Shared Resource is supported by the Vanderbilt Ingram Cancer Center (P30 CA68485) and the Vanderbilt Digestive Disease Research Center (DK058404).

For work described in this manuscript, I.S.G., A.R.S., K.J.K., S.W., K.A.P., R.V., N.R., E.F.F., C.M.H. were supported in part by NIH NIAID award R01AI131722-S1, the Hays Foundation COVID-19 Research Fund, and Fast Grants. J.S.M and D.W. were supported in part by a National Institutes of Health (NIH)/National Institute of Allergy and Infectious Diseases (NIAID) grant awarded to J.S.M. (R01-AI127521). L.M. and S.I.R. acknowledge research funding from the South African Medical Research Council (MRC) Extramural Unit and SHIP programs and an H3 Africa grant (U01A136677). S.I.R. is supported by the South African Research Chairs Initiative of the Department of Science and Technology and the NRF (Grant No 98341). R.B., A.S., D.R.M., were supported by NIH grants (U54CA260543, R01AI157155). P.A. and K.J. were supported by NIH grant R01 AI14567. J.E.C., R.H.C., N.S., R.N.S., and R.E.S., were supported by Defense Advanced Research Projects Agency (DARPA) grants HR0011-18-2-0001 and HR00 11-18-3-0001; NIH contracts 75N93019C00074 and 75N93019C00062; NIH grants U01 AI150739, R01 AI130591 and R35 HL145242; the Dolly Parton COVID-19 Research Fund at Vanderbilt; and NIH grant S10 RR028106 for the Next Generation Nucleic Acid Sequencer, housed in VANTAGE.M.S.D. and R.E.C. were supported by grants from NIH (R01 AI157155) and the Defense Advanced Research Project Agency (HR001117S0019). B.F.H. and R.P. were supported by NC State funding for COVID research. B.S.G. was supported by intramural funding from the NIAID. C.M.H. was supported in part by NIH grant T32 GM008320-30. D.R.M. was supported by an NIH F32 AI152296, a Burroughs Wellcome Fund Postdoctoral Enrichment Program Award, and was previously supported by an NIH NIAID T32 AI007151.

## METHODS

### Donor Information

A donor was identified and enrolled in VRC 200 sample collection clinical trial #NCT00067054 at the NIH Clinical Center and samples were collected following informed consent. The protocol was approved by the NIAID Institutional Review Board and all applicable human subject’s protections research requirements were followed. The donor had prior SARS-CoV-1 infection during the 2004 outbreak in Hong Kong, and the PBMC sample was collected over 10 years post infection (20 million PBMCs).

### Antigen Purification

A variety of recombinant soluble protein antigens were used in the LIBRA-seq experiment and other experimental assays.

Plasmids encoding residues 1–1208 of the SARS-CoV-2 spike with a mutated S1/S2 cleavage site, proline substitutions at positions 986 and 987, and a C-terminal T4-fibritin trimerization motif, an 8x HisTag, and a TwinStrepTag (SARS-CoV-2 S-2P); residues 1-1190 of the SARS-CoV-1 spike with proline substitutions at positions 968 and 969, and a C-terminal T4-fibritin trimerization motif, an 8x HisTag, and a TwinStrepTag (SARS-CoV-1 S-2P); residues 1-1291 of the MERS-CoV spike with a mutated S1/S2 cleavage site, proline substitutions at positions 1060 and 1061, and a C-terminal T4-fibritin trimerization motif, an AviTag, an 8x HisTag, and a TwinStrepTag (MERS-CoV S-2P Avi); residues 1-751 of the MERS-CoV spike with a C-terminal T4-fibritin trimerization motif, 8x HisTag, and a TwinStrepTag (MERS-CoV S1); residues 1-1277 of the HCoV-HKU1 spike with a mutated S1/S2 cleavage site, proline substitutions at positions 1067 and 1068, and a C-terminal T4-fibritin trimerization motif, an 8x HisTag, and a TwinStrepTag (HCoV-HKU1 S-2P); residues 1-1278 of the HCoV-OC43 spike with proline substitutions at positions 1070 and 1071, and a C-terminal T4-fibritin trimerization motif, an 8x HisTag, and a TwinStrepTag (HCoV-OC43 S-2P); or residues 319–591 of SARS-CoV-2 S with a C-terminal monomeric human IgG Fc-tag and an 8x HisTag (SARS-CoV-2 RBD-SD1) were transiently transfected into FreeStyle293F cells (Thermo Fisher) using polyethylenimine. The coronavirus trimer spike antigens were in a prefusion-stabilized (S-2P) conformation that better represents neutralization-sensitive epitopes in comparison to their wild-type forms^37^. Two hours post-transfection, cells were treated with kifunensine to ensure uniform glycosylation. Transfected supernatants were harvested after 6 days of expression. SARS-CoV-2 RBD-SD1 was purified using Protein A resin (Pierce), SARS-CoV-2 S-2P, SARS-CoV-1 S-2P, MERS-CoV S-2P Avi, MERS-CoV S1, HCoV-HKU1 S-2P and HCoV-OC43 S-2P were purified using StrepTactin resin (IBA). Affinity-purified SARS-CoV-2 RBD-SD1 was further purified over a Superdex75 column (GE Life Sciences). MERS-CoV S1 was purified over a Superdex200 Increase column (GE Life Sciences). SARS-CoV-2 S-2P, SARS-CoV-1 S-2P, MERS-CoV S-2P Avi, HCoV-HKU1 S-2P and HCoV-OC43 S-2P were purified over a Superose6 Increase column (GE Life Sciences).

For the HIV-1 gp140 SOSIP variant from strain ZM197 (clade C) and CZA97 (clade C), recombinant, soluble antigens contained an AviTag and were expressed in Expi293F cells using polyethylenimine transfection reagent and cultured. FreeStyle F17 expression medium supplemented with pluronic acid and glutamine was used. The cells were cultured at 37°C with 8% CO_2_ saturation and shaking. After 5-7 days, cultures were centrifuged and supernatant was filtered and run over an affinity column of agarose bound *Galanthus nivalis* lectin. The column was washed with PBS and antigens were eluted with 30 mL of 1M methyl-a-D-mannopyranoside. Protein elutions were buffer exchanged into PBS, concentrated, and run on a Superdex 200 Increase 10/300 GL Sizing column on the AKTA FPLC system. Fractions corresponding to correctly folded protein were collected, analyzed by SDS-PAGE and antigenicity was characterized by ELISA using known monoclonal antibodies specific to each antigen. Avitagged antigens were biotinylated using BirA biotin ligase (Avidity LLC).

For binding studies, SARS-CoV-2 HexaPro S, SARS-CoV-1 S, SARS-CoV-2 RBD, SARS-CoV-1 RBD, and MERS-CoV RBD constructs were expressed in the transient expression system previously mentioned. S proteins were purified using StrepTrap HP columns and RBD constructs were purified over protein A resin, respectively. Each resulting protein was further purified to homogeneity by size-exclusion chromatography on a Superose 6 10/300 GL column.

SARS-CoV-2 S1, SARS-CoV-2 S1 D614G, SARS-CoV-2 S2, and SARS-CoV-2 NTD truncated proteins were purchased from the commercial vendor, Sino Biological.

### DNA-barcoding of Antigens

We used oligos that possess 15 bp antigen barcode, a sequence capable of annealing to the template switch oligo that is part of the 10X bead-delivered oligos, and contain truncated TruSeq small RNA read 1 sequences in the following structure: 5’-CCTTGGCACCCGAGAATTCCANNNNNNNNNNNNNCCCATATAAGA*A*A-3’, where Ns represent the antigen barcode. We used the following antigen barcodes: GCTCCTTTACACGTA (SARS-CoV-2 S), TGACCTTCCTCTCCT (SARS-CoV-1 S), ACAATTTGTCTGCGA (MERS-CoV S), TCCTTTCCTGATAGG (MERS-CoV S1), CAGGTCCCTTATTTC (HKU1-CoV S), TAACTCAGGGCCTAT (OC43-CoV S), CAGCCCACTGCAATA (CZA97), and ATCGTCGAGAGCTAG (ZM197). Oligos were ordered from IDT with a 5’ amino modification and HPLC purified.

For each antigen, a unique DNA barcode was directly conjugated to the antigen itself. In particular, 5’amino-oligonucleotides were conjugated directly to each antigen using the Solulink Protein-Oligonucleotide Conjugation Kit (TriLink cat no. S-9011) according to manufacturer’s instructions. Briefly, the oligo and protein were desalted, and then the amino-oligo was modified with the 4FB crosslinker, and the biotinylated antigen protein was modified with S-HyNic. Then, the 4FB-oligo and the HyNic-antigen were mixed together. This causes a stable bond to form between the protein and the oligonucleotide. The concentration of the antigen-oligo conjugates was determined by a BCA assay, and the HyNic molar substitution ratio of the antigen-oligo conjugates was analyzed using the NanoDrop according to the Solulink protocol guidelines. AKTA FPLC was used to remove excess oligonucleotide from the protein-oligo conjugates, which were also verified using SDS-PAGE with a silver stain. Antigen-oligo conjugates were also used in flow cytometry titration experiments.

### Antigen specific B cell sorting

Cells were stained and mixed with DNA-barcoded antigens and other antibodies, and then sorted using fluorescence activated cell sorting (FACS). First, cells were counted and viability was assessed using Trypan Blue. Then, cells were washed three times with DPBS supplemented with 0.1% Bovine serum albumin (BSA). Cells were resuspended in DPBS-BSA and stained with cell markers including viability dye (Ghost Red 780), CD14-APC-Cy7, CD3-FITC, CD19-BV711, and IgG-PE-Cy5. Additionally, antigen-oligo conjugates were added to the stain. After staining in the dark for 30 minutes at room temperature, cells were washed three times with DPBS-BSA at 300 g for five minutes. Cells were then incubated for 15 minutes at room temperature with Streptavidin-PE to label cells with bound antigen. Cells were washed three times with DPBS-BSA, resuspended in DPBS, and sorted by FACS. Antigen positive cells were bulk sorted and delivered to the Vanderbilt Technologies for Advanced Genomics (VANTAGE) sequencing core at an appropriate target concentration for 10X Genomics library preparation and subsequent sequencing. FACS data were analyzed using FlowJo.

### Sample preparation, library preparation, and sequencing

Single-cell suspensions were loaded onto the Chromium Controller microfluidics device (10X Genomics) and processed using the B-cell Single Cell V(D)J solution according to manufacturer’s suggestions for a target capture of 10,000 B cells per 1/8 10X cassette, with minor modifications in order to intercept, amplify and purify the antigen barcode libraries as previously described^24^.

### Sequence processing and bioinformatic analysis

We utilized and modified our previously described pipeline to use paired-end FASTQ files of oligo libraries as input, process and annotate reads for cell barcode, UMI, and antigen barcode, and generate a cell barcode - antigen barcode UMI count matrix^24^. BCR contigs were processed using Cell Ranger (10X Genomics) using GRCh38 as reference. Antigen barcode libraries were also processed using Cell Ranger (10X Genomics). The overlapping cell barcodes between the two libraries were used as the basis of the subsequent analysis. We removed cell barcodes that had only non-functional heavy chain sequences as well as cells with multiple functional heavy chain sequences and/or multiple functional light chain sequences, reasoning that these may be multiplets. Additionally, we aligned the BCR contigs (filtered_contigs.fasta file output by Cell Ranger, 10X Genomics) to IMGT reference genes using HighV-Quest^38^. The output of HighV-Quest was parsed using ChangeO^39^ and merged with an antigen barcode UMI count matrix. Finally, we determined the LIBRA-seq score for each antigen in the library for every cell as previously described^24^.

### Antibody Expression and Purification

For each antibody, variable genes were inserted into custom plasmids encoding the constant region for the IgG1 heavy chain as well as respective lambda and kappa light chains (pTwist CMV BetaGlobin WPRE Neo vector, Twist Bioscience). Antibodies were expressed in Expi293F mammalian cells (ThermoFisher) by co-transfecting heavy chain and light chain expressing plasmids using polyethylenimine transfection reagent and cultured for 5-7 days. Cells were maintained in FreeStyle F17 expression medium supplemented at final concentrations of 0.1% Pluronic Acid F-68 and 20% 4mM L-Glutamine. These cells were cultured at 37°C with 8% CO_2_ saturation and shaking. After transfection and 5-7 days of culture, cell cultures were centrifuged and supernatant was 0.45 μm filtered with Nalgene Rapid Flow Disposable Filter Units with PES membrane. Filtered supernatant was run over a column containing Protein A agarose resin equilibrated with PBS. The column was washed with PBS, and then antibodies were eluted with 100 mM Glycine HCl at 2.7 pH directly into a 1:10 volume of 1M Tris-HCl pH 8.0. Eluted antibodies were buffer exchanged into PBS 3 times using Amicon Ultra centrifugal filter units and concentrated. Antibodies were analyzed by SDS-PAGE. Additionally, antibodies 46472-1, 46472-2, 46472-3, 46472-4, 46472-6 and 46472-12 were assessed by size exclusion chromatography on a Superdex 200 Increase 10/300 GL Sizing column with the AKTA FPLC system.

### High-throughput Antibody Expression

For high-throughput production of recombinant antibodies, approaches were used that are designated as microscale. For antibody expression, microscale transfection were performed (~1 ml per antibody) of CHO cell cultures using the Gibco ExpiCHO Expression System and a protocol for deep 96-well blocks (Thermo Fisher Scientific). In brief, synthesized antibody-encoding DNA (~2 μg per transfection) was added to OptiPro serum free medium (OptiPro SFM), incubated with ExpiFectamine CHO Reagent and added to 800 μl of ExpiCHO cell cultures into 96-deep-well blocks using a ViaFlo 384 liquid handler (Integra Biosciences). The plates were incubated on an orbital shaker at 1,000 r.p.m. with an orbital diameter of 3 mm at 37 °C in 8% CO_2_. The next day after transfection, ExpiFectamine CHO Enhancer and ExpiCHO Feed reagents (Thermo Fisher Scientific) were added to the cells, followed by 4 d incubation for a total of 5 d at 37 °C in 8% CO_2_. Culture supernatants were collected after centrifuging the blocks at 450*g* for 5 min and were stored at 4°C until use. For high-throughput microscale antibody purification, fritted deep-well plates were used containing 25 μl of settled protein G resin (GE Healthcare Life Sciences) per well. Clarified culture supernatants were incubated with protein G resin for mAb capturing, washed with PBS using a 96-well plate manifold base (Qiagen) connected to the vacuum and eluted into 96-well PCR plates using 86 μl of 0.1 M glycine-HCL buffer pH 2.7. After neutralization with 14 μl of 1 M Tris-HCl pH 8.0, purified mAbs were buffer-exchanged into PBS using Zeba Spin Desalting Plates (Thermo Fisher Scientific) and stored at 4°C until use.

### ELISA

To assess antibody binding, soluble protein was plated at 2 μg/ml overnight at 4°C. The next day, plates were washed three times with PBS supplemented with 0.05% Tween-20 (PBS-T) and coated with 5% milk powder in PBS-T. Plates were incubated for one hour at room temperature and then washed three times with PBS-T. Primary antibodies were diluted in 1% milk in PBS-T, starting at 10 μg/ml with a serial 1:5 dilution and then added to the plate. The plates were incubated at room temperature for one hour and then washed three times in PBS-T. The secondary antibody, goat anti-human IgG conjugated to peroxidase, was added at 1:10,000 dilution in 1% milk in PBS-T to the plates, which were incubated for one hour at room temperature. Goat anti-mouse secondary was used for SARS-CoV-1 specific control antibody 240CD (BEI Resources). Plates were washed three times with PBS-T and then developed by adding TMB substrate to each well. The plates were incubated at room temperature for ten minutes, and then 1N sulfuric acid was added to stop the reaction. Plates were read at 450 nm. Data are represented as mean ± SEM for one ELISA experiment. ELISAs were repeated 2 or more times. The area under the curve (AUC) was calculated using GraphPad Prism 8.0.0. For antibody 240CD, the following reagent was obtained through BEI Resources, NIAID, NIH: Monoclonal Anti-SARS-CoV S Protein (Similar to 240C), NR-616.

### Competition ELISA

Competition ELISAs were performed as described above, with some modifications. After coating with antigen and blocking, 25 μl of non-biotinylated competitor antibody was added to each well at 10 μg/ml and incubated at 37°C for 10 minutes. Then, without washing, 75 μl biotinylated antibody (final concentration of 1 μg/ml) was added and incubated at 37°C for 1 hour. After washing three times with PBS-T, streptavidin-HRP was added at 1:10,000 dilution in 1% milk in PBS-T and incubated for 1 hour at room temperature. Plates were washed and substrate and sulfuric acid were added as described above. ELISAs were repeated at least 2 times. Data is shown as the % decrease in binding.

### Autoreactivity

Monoclonal antibody reactivity to nine autoantigens (SSA/Ro, SS-B/La, Sm, ribonucleoprotein (RNP), Scl 70, Jo-1, dsDNA, centromere B, and histone) was measured using the AtheNA Multi-Lyte^®^ ANA-II Plus test kit (Zeus scientific, Inc, #A21101). Antibodies were incubated with AtheNA beads for 30min at concentrations of 50, 25, 12.5 and 6.25 μg/mL. Beads were washed, incubated with secondary and read on the Luminex platform as specified in the kit protocol. Data were analyzed using AtheNA software. Positive (+) specimens received a score >120, and negative (-) specimens received a score <100. Samples between 100-120 were considered indeterminate.

### Mannose competition

Mannose competition ELISAs were performed as described above with minor modifications. After antigen coating and washing, nonspecific binding was blocked by incubation with 5% FBS diluted in PBS for 1 hour at RT. Primary antibodies were diluted in 5% FBS-PBST +/- 1M D-(+)- Mannose starting at 10 μg/ml with a serial 1:5 dilution and then added to the plate for 1 hour at RT. After washing, antibody binding was detected with goat anti-human IgG conjugated to peroxidase and added at 1:10,000 dilution in 5% FBS in PBS-T to the plates. After 1 hour incubation, plates were washed and substrate and sulfuric acid were added as described above. Data shown is representative of three replicates.

### Epitope Mapping Visualization

SARS-CoV-2 Spike (PDB-6VSB) was visualized using PyMOL software. Antibody epitopes were visualized on the SARS-CoV-2 spike using a structure of the pre-fusion stabilized SARS-CoV-2 S-2P construct^5^ modeled in the molecular graphics software PyMOL (The PyMOL Molecular Graphics System, Version 2.3.5 Schrödinger, LLC).

### RTCA Neutralization Assay

To assess for neutralizing activity against SARS-CoV-2 strain 2019 n-CoV/USA_WA1/2020 (obtained from the Centers for Disease Control and Prevention, a gift from N. Thornburg), we used the high-throughput RTCA assay and xCelligence RTCA HT Analyzer (ACEA Biosciences) that has been described previously^11^. After obtaining a background reading of a 384-well E-plate, 6,000 Vero-furin cells^40^ were seeded per well. Sensograms were visualized using RTCA HT software version 1.0.1 (ACEA Biosciences). One day later, equal volumes of virus were added to antibody samples and incubated for 1 h at 37°C in 5% CO_2_. mAbs were tested in triplicate with a single (1:20) dilution. Virus–mAb mixtures were then added to Vero-furin cells in 384-well E-plates. Controls were included that had Vero-furin cells with virus only (no mAb) and media only (no virus or mAb). E-plates were read every 8-12 h for 72 h to monitor virus neutralization. At 32 h after virus-mAb mixtures were added to the E-plates, cell index values of antibody samples were compared to those of virus only and media only to determine presence of neutralization.

### Nano-luciferase Neutralization Assay

A full-length SARS-CoV-2 virus based on the Seattle Washington isolate and a full-length SARS-CoV virus based on the Urbani isolate were designed to express luciferase and was recovered via reverse genetics and described previously^41,42^. Viruses were titered in Vero E6 USAMRID cells to obtain a relative light units (RLU) signal of at least 10X the cell only control background. Vero E6 USAMRID cells were plated at 20,000 cells per well the day prior in clear bottom black walled 96-well plates (Corning 3904). Neutralizing antibody serum samples were tested at a starting dilution of 1:40 and were serially diluted 4-fold up to eight dilution spots. Antibody-virus complexes were incubated at 37C with 5% CO2 for 1 hour. Following incubation, growth media was removed and virus-antibody dilution complexes were added to the cells in duplicate. Virus-only controls and cell-only controls were included in each neutralization assay plate. Following infection, plates were incubated at 37C with 5% CO2 for 48 hours. After the 48 hour incubation, cells were lysed and luciferase activity was measured via Nano-Glo Luciferase Assay System (Promega) according to the manufacturer specifications. SARS-CoV and SARS-CoV-2 neutralization titers were defined as the sample dilution at which a 50% reduction in RLU was observed relative to the average of the virus control wells.

### SPR

His-tagged SARS-CoV-2 RBD-SD1 was immobilized to a NiNTA sensorchip to a level of ~150 RUs using a Biacore X100. Serial dilutions of purified Fab 46472-12 were evaluated for binding, ranging in concentration from 1 to 0.25 μM. The resulting data were fit to a 1:1 binding model using Biacore Evaluation Software.

### Fc Effector function Assays

#### Antibody-dependent Cellular Phagocytosis (ADCP)

Antibody-dependent cellular phagocytosis (ADCP) was performed using biotinylated SARS-CoV-2 or SARS-CoV-1 S coated fluorescent neutravidin beads as previously described^43^. Briefly, beads were incubated for two hours with antibodies at a starting concentration of 50μg/ml and titrated five fold. CR3022 was used as a positive control while Palivizumab was used as a negative control. Antibodies and beads were incubated with THP-1 cells overnight, fixed and interrogated on the FACSAria II. Phagocytosis score was calculated as the percentage of THP-1 cells that engulfed fluorescent beads multiplied by the geometric mean fluorescence intensity of the population in the FITC channel less the no antibody control.

#### Antibody-dependent Cellular Trogocytosis (ADCT)

ADCT was performed as described in and modified from a previously described study^26^. HEK293T-ACE2 expressing cells were pulsed with SARS-CoV-2 S protein (10μg/ml) for 75 minutes or HEK293T cells transfected with a SARS-CoV-2 spike pcDNA vector were surface biotinylated with EZ-Link Sulfo-NHS-LC-Biotin as recommended by the manufacturer. Fifty-thousand cells per well were incubated with antibody for 30 minutes starting at 25μg/ml and titrated 5 fold. CR3022 was used as a positive control with Palivizumab as a negative. Following a RPMI media wash, these were then incubated with carboxyfluorescein succinimidyl ester (CFSE) stained THP-1 cells (5 X10^4^ cells per well) for 1 hour and washed with 15mM EDTA/PBS followed by PBS. Cells were then stained for biotin using Streptavidin-PE and read on a FACSAria II. Trogocytosis score was determined as the proportion of CFSE positive THP-1 cells also positive for streptavidin-PE less the no antibody control.

#### Antibody-dependent Complement Deposition (ADCD)

Antibody-dependent complement deposition was performed as previously described^44^. Briefly biotinylated SARS-Cov-2 S protein was coated 1:1 onto fluorescent neutravidin beads for 2 hours at 37 degrees. These beads were incubated with 100ug/ml of antibody for 1 hour and incubated with guinea pig complement diluted 1 in 50 with gelatin/veronal buffer for 15 minutes at 37 degrees. Beads were washed at 2000g twice in PBS and stained with guinea pig C3b-FITC, fixed and interrogated on a FACSAria II. Complement deposition score was calculated as the percentage of C3b-FITC positive beads multiplied by the geometric mean fluorescent intensity of FITC in this population less the no antibody or heat inactivated controls.

#### Antibody Prophylaxis - Murine Model of Infection

12-month old female BALB/c mice (BALB/cAnHsd; Envigo, stock number 047) were treated with 200 μg mAb intraperitoneally (i.p.) 12 hours prior to virus inoculation. The next day, mice were administered intranasally with 1×10^3^ PFU or 1×10^4^PFU of SARS-CoV-2 MA10, respectively. Mice were monitored daily for weight loss, morbidity, and mortality, and after four days one lung lobe was taken for pathological analysis and the other lobe was processed for qPCR and viral load determination as previously described^28^. Gross pulmonary hemorrhage was observed at time of tissue harvest and scored on a scale of 0 (no hemorrhage in any lobe, normal pink healthy lung) to 4 (complete hemorrhage in all lobes of the lung, completely dark red lung).

For viral titer and hemorrhage score comparisons, an ordinary one-way ANOVA test with multiple comparisons was performed using Prism software, GraphPad Prism version 8.0.

#### ACE2 Binding Inhibition Assay

Wells of 384-well microtiter plates were coated with purified recombinant SARS-CoV-2 S-2P ectoprotein at 4°C overnight. Plates were blocked with 2% non-fat dry milk and 2% normal goat serum in DPBS-T for 1 hr. Purified mAbs were diluted two-fold in blocking buffer starting from 10 μg/mL in triplicate, added to the wells (20 μL/well), and incubated at ambient temperature. Recombinant human ACE2 with a C-terminal FLAG tag protein was added to wells at 2 μg/mL in a 5 μL/well volume (final 0.4 μg/mL concentration of ACE2) without washing of antibody and then incubated for 40 min at ambient temperature. Plates were washed, and bound ACE2 was detected using HRP-conjugated anti-FLAG antibody and TMB substrate. ACE2 binding without antibody served as a control. The signal obtained for binding of the ACE2 in the presence of each dilution of tested antibody was expressed as a percentage of the ACE2 binding without antibody after subtracting the background signal.

#### Computational Identification of Residue-level Knockout Mutants

Potential cross-reactive epitopes were identified based on sequence and structural homology. Reference sequences for each Coronavirus S were obtained either from NCBI for SARS-CoV-2 (YP_009724390.1) and MERS-CoV (YP_009047204.1) or from Uniprot for SARS-CoV-1 (P59594) of the spikes was then obtained using MUSCLE^45^ and the amino acid similarity to SARS-CoV-2 at each residue position was calculated using the BLOSUM-62 scoring matrix^46^. These scores were then used to color each residue position on the SARS-CoV-2 S structure (PDB ID: 6VSB) in PyMOL (Schrodinger, version 2.3.5) in order to visualize surface patches and linear epitopes with structural homology. These conserved regions were then visualized on the other human coronavirus spike structures by retrieving them from the Protein Databank (SARS-CoV-1: 5X5B, MERS-CoV: 5W9I) and aligning them to the SARS-CoV-2 S structure. Finally, the residue N165 was part of a conserved surface patches and was mutated to alanine and tested for binding with antibodies. The N709A mutant tested was previously described in Acharya et al., BioRxiv (2020).

## QUANTIFICATION AND STATISTICAL ANALYSIS

ELISA error bars (standard error of the mean) were calculated using GraphPad Prism version 8.0.0. ANOVA analysis was performed on viral load titers and hemorrhage scores from animal experiments using GraphPad Prism version 8.0.0.

