## Supplemental for "Cross-reactive coronavirus antibodies with diverse epitope specificities and extra-neutralization functions"

#### Supplemental Materials

#### Supplemental Figure Captions

**Figure S1.**

(**A**) Gating scheme for fluorescent-activated cell sorting of convalescent SARS-CoV-1 donor. Cells were initially stained with Ghost Red 780, CD14-APC-Cy7, CD3-FITC, CD19-BV711, and IgG-PE-Cy5 along with a DNA-barcoded antigen screening library. To detect antigen-positive B cells, cells were washed and treated with a streptavidin-PE secondary stain. Gates as drawn are based on gates used during the sort, and percentages from the sort are listed.

(**B**) The categorization of processing of Cell Ranger identified cells after sequencing is shown.

(**C**) Genetic sequence characteristics of 15 antibody candidates. Percent identity is calculated at the nucleotide level and CDRH3 and CDRL3 lengths and sequences are noted at the amino acid level.

(**D**) ELISA binding data against coronavirus S antigens. HIV-specific antibody VRC01 was used as a negative control and anti-SARS-CoV-1 mouse antibody 240CD was used as a positive control (BEI Resources). ELISAs were performed in technical duplicates with at least two biological duplicates.

**Figure S2.**

(**A**) Cross-reactive antibodies were tested for binding to SARS-CoV-2 S1 domain, SARS-CoV-2 S1 domain D614G, SARS-CoV-2 S2 domain, and SARS-CoV-2 S (HexaPro). Anti-HIV antibody VRC01 is shown as a negative control and anti-SARS-CoV-1 antibody 240CD is shown as positive control.

(**B**) S1-directed antibodies 46472-6 and 46472-12 were tested for binding against SARS-CoV-2 RBD, SARS-CoV-1 RBD, SARS-CoV-2 NTD, and SARS-CoV-2 S (HexaPro). Anti-HIV antibody

VRC01 is shown as a negative control and anti-SARS-CoV-1 antibody 240CD is shown as positive control.

(**C**) 46472-12 was tested for its ability to block ACE2 binding to SARS-CoV-2 S. Signal shown is anti-Flag tag detection of an ACE2-Flag tag protein construct.

(**D**) 46472-6 and 46472-12 were tested for binding to SARS-CoV-2 S (HexaPro) mutants, N165A and N709A, by ELISA.

(**E**) Mannose competition binding assays were performed to see if cross-reactive antibody binding to SARS-CoV-2 S could be modulated by mannose.

**Figure S3.**

(**A**) Antibodies were tested for neutralization in a SARS-CoV-1 and SARS-CoV-2 nano-luciferase neutralization assay.

(**B**) Antibodies were tested for neutralization in a SARS-CoV-2 RTCA assay.

(**C**) Cross-reactive coronavirus antibodies were tested for ability to mediate antibody-dependent cellular phagocytosis against SARS-CoV-2 S.

(**D**) Cross-reactive coronavirus antibodies were tested for ability to mediate antibody-dependent cellular phagocytosis against SARS-CoV-1 S coated on beads.

(**E**) Cross-reactive coronavirus antibodies were tested for ability to mediate antibody-dependent cellular trogocytosis against SARS-CoV-2 S coated on cells.

(**F**) Cross-reactive coronavirus antibodies were tested for ability to mediate antibody-dependent cellular trogocytosis against transfected cells displaying SARS-CoV-2 S

(**G**) Cross-reactive coronavirus antibodies were tested for ability to mediate antibody-dependent complement deposition against SARS-CoV-2 S.

**Figure S4.**

(**A**) For each antibody treatment group for the experiment utilizing $1x10^3$ PFU of SARS-CoV-2 MA, a table showing the number of animals to survive per group, per day is shown. Body weights of each mouse in the four treatment groups were measured daily.

(**B**) RT-qPCR quantification of lung viral titer is shown.

(**C**) For each antibody treatment group for the experiment utilizing $1x10^4$ PFU of SARS-CoV-2 MA, a table showing the number of animals to survive per group, per day is shown (survival curves shown in **Figure 4C**). 2/5, 4/5, 3/5, and 2/5 mice survived to day 4 for antibodies 46472-4, 46472-12, CR3022 and isotype control DENV-2D22 respectively. Body weights of each mouse in the four treatment groups in both experiments were measured daily.

(**D**) RT-qPCR quantification of lung viral burden is shown.

### Supplemental Figure 1

A

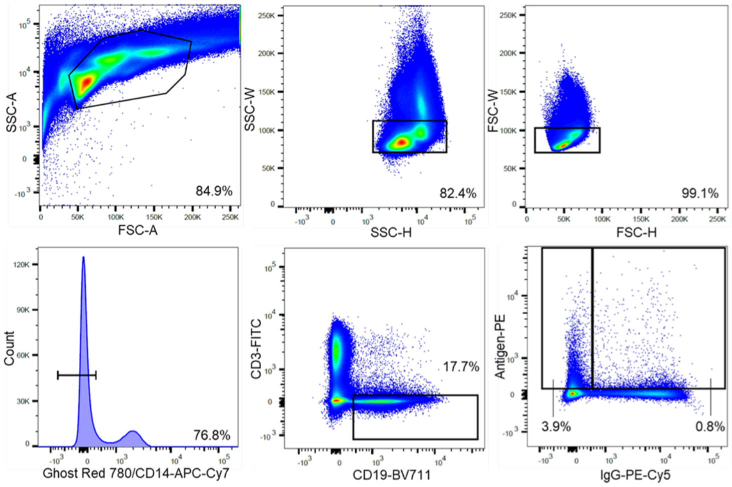


B

| | 4647-2 Processing |
|---|---|
| Unique Heavy Chain Barcodes | 5113 |
| Unique Heavy Chain Barcodes With Functional Heavy Chains | 5072 |
| Unique Heavy Chain Barcodes Without Cells With Multiple Heavy Chains | 4875 |
| Unique Light Chain Barcodes | 7593 |
| Unique Light Chain Barcodes With Functional Light Chains | 7586 |
| Unique Light Chain Barcodes Without Cells With Multiple Light Chains | 5914 |
| Unique Paired Heavy-Light Chain Barcodes | 3813 |
| Overlapping Barcodes Between Paired Heavy-Light Chains and Antigen Counts | 2625 |

C

| Name | VH gene | VH % identity | CDRH3 length | VDJ junction | VL gene | VL % identity | CDRL3 length | VJ junction |
|---|---|---|---|---|---|---|---|---|
| **46472_1** | IGHV1-69 | 93.75 | 12 | AREVNYYSAFDD | IGKV3-15 | 97.85 | 8 | QQYNFWWT |
| **46472_2** | IGHV5-10 | 94.10 | 20 | AAGPTGYDLLTGQYFPYFNY | IGLV6-57 | 95.19 | 9 | QSYHGSDVV |
| **46472_3** | IGHV3-30 | 94.79 | 14 | ARDRSATYYGPFDY | IGLV3-19 | 94.98 | 12 | NSRDNSGNHPVI |
| **46472_4** | IGHV3-23 | 90.97 | 19 | AKDLLSHSGTYSAGSTFDY | IGKV1-39 | 94.98 | 8 | QESYSTNT |
| **46472_5** | IGHV4-39 | 84.19 | 9 | ARLDYSKQT | IGKV2-28 | 96.26 | 9 | MQALQTPLT |
| **46472_6** | IGHV4-39 | 90.72 | 24 | ARRPQYLLLSMTTGRRHHDFVMDV | IGKV4-1 | 97.31 | 9 | QQYYNTPRT |
| **46472_7** | IGHV3-64 | 92.36 | 11 | VKEDTPLVFDS | IGKV1-5 | 93.19 | 8 | QQYDSYST |
| **46472_8** | IGHV3-53 | 84.91 | 12 | ALGRKDYGDYYR | IGKV3-20 | 89.01 | 11 | QQYAPSPPWYI |
| **46472_9** | IGHV4-61 | 95.53 | 15 | ASLPTYGSGRWGIDS | IGKV1-12 | 96.42 | 9 | QQGNSFPLT |
| **46472_10** | IGHV3-53 | 92.63 | 16 | AGFLPVYNNGWSYFDS | IGLV1-44 | 95.09 | 11 | AVWDDSLNGPV |
| **46472_11** | IGHV3-23 | 87.15 | 13 | VKMRTAVVGVTPL | IGKV1-6 | 95.34 | 9 | LQDYNYLFS |
| **46472_12** | IGHV1-8 | 92.01 | 20 | ARDRVERTGNVGFGYYAMDV | IGKV1-39 | 95.34 | 10 | QQTYSSPSYT |
| **46472_13** | IGHV3-7 | 90.97 | 15 | ARVTIVSSFTNRFDP | IGKV1-9 | 90.32 | 5 | QHRVT |
| **46472_14** | IGHV3-53 | 82.81 | 6 | VRGRTY | IGKV3-15 | 92.47 | 9 | QQYNRWLWT |
| **46472_15** | IGHV7-4 | 97.57 | 23 | ARDFDLVVPSATYPPFYYHGMDV | IGLV3-19 | 97.49 | 13 | NSRDSSGDQTFYV |

D

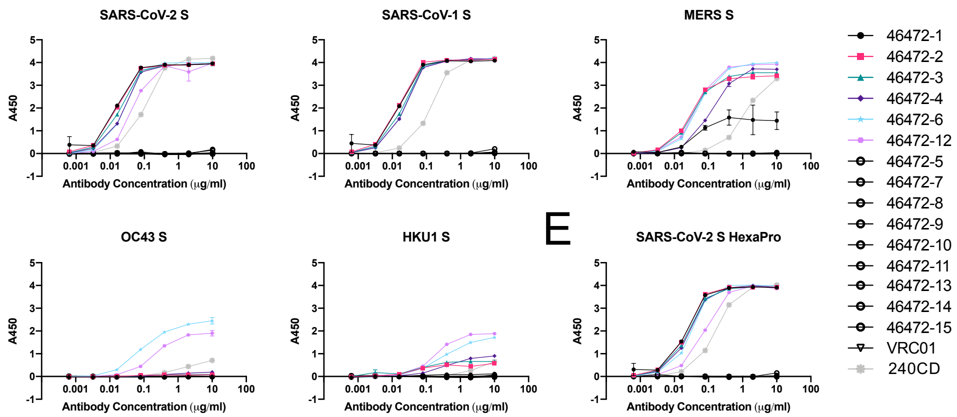

### Supplemental Figure 2

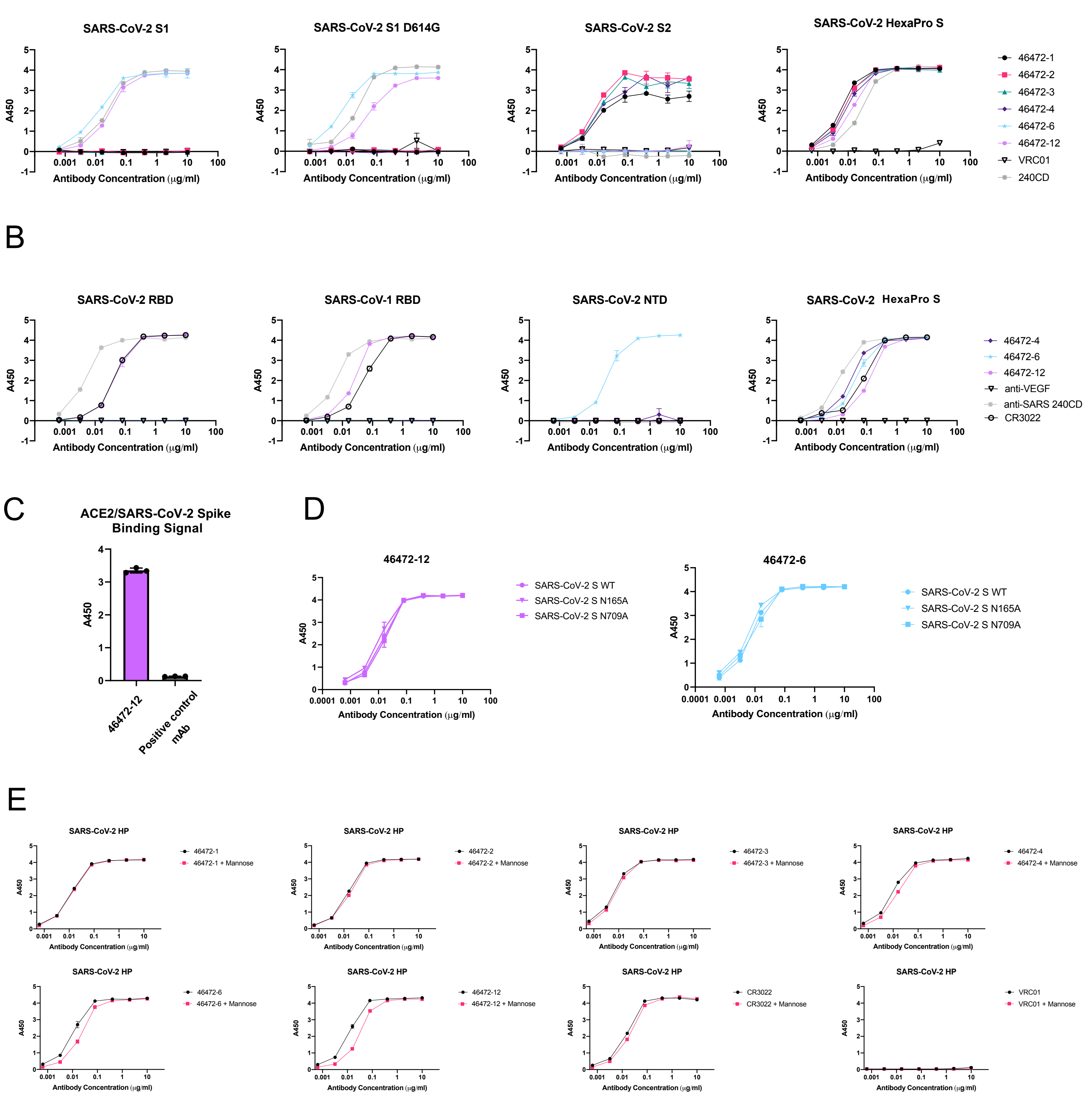

Supplemental Figure 3

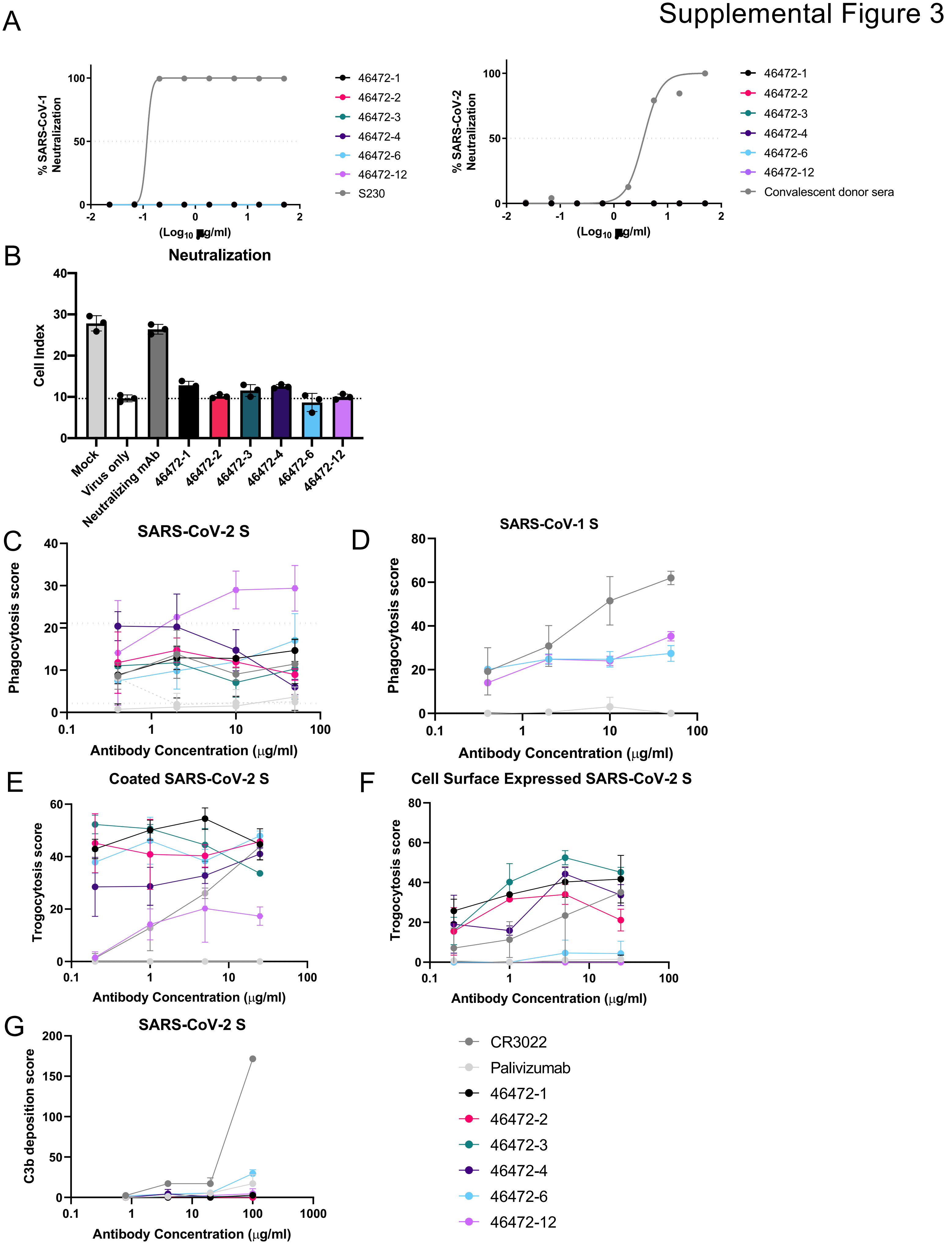

Supplemental Figure 4

A

| Antibody | Animal Survival, Days Post Infection | | | | |
|---|---|---|---|---|---|
| | 0 | 1 | 2 | 3 | 4 |
| **46472-12** | 4/4 | 4/4 | 4/4 | 4/4 | 4/4 |
| **46472-4** | 5/5 | 5/5 | 5/5 | 5/5 | 5/5 |
| **CR3022** | 5/5 | 5/5 | 5/5 | 5/5 | 5/5 |
| **DENV-2D22** | 5/5 | 5/5 | 5/5 | 5/5 | 4/5 |

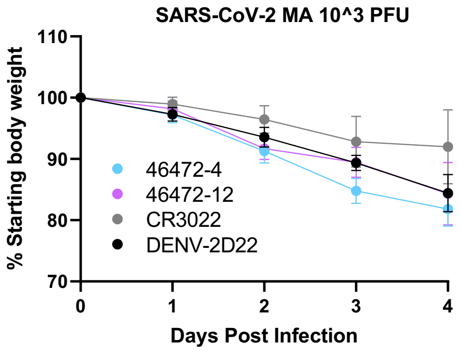


B

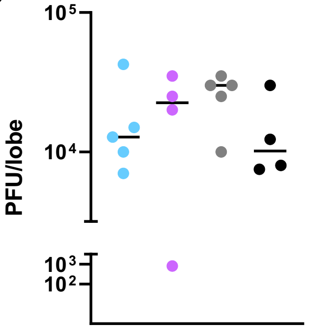


C

| Antibody | Animal Survival, Days Post Infection | | | | |
|---|---|---|---|---|---|
| | 0 | 1 | 2 | 3 | 4 |
| **46472-12** | 5/5 | 5/5 | 5/5 | 5/5 | 4/5 |
| **46472-4** | 5/5 | 5/5 | 5/5 | 4/5 | 2/5 |
| **CR3022** | 5/5 | 5/5 | 5/5 | 4/5 | 3/5 |
| **DENV-2D22** | 5/5 | 5/5 | 5/5 | 5/5 | 2/5 |

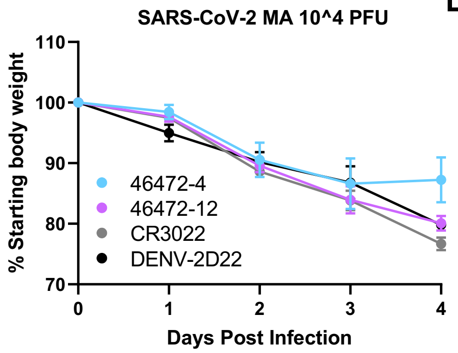


D

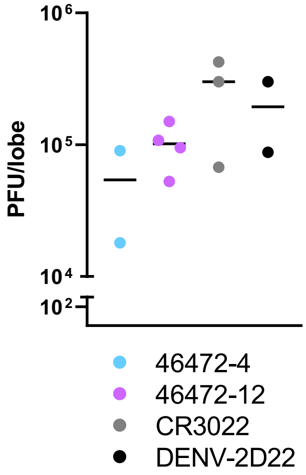
